# Inference under a Wright-Fisher model using an accurate beta approximation

**DOI:** 10.1101/021261

**Authors:** Paula Tataru, Thomas Bataillon, Asger Hobolth

## Abstract

The large amount and high quality of genomic data available today enables, in principle, accurate inference of evolutionary history of observed populations. The Wright-Fisher model is one of the most widely used models for this purpose. It describes the stochastic behavior in time of allele frequencies and the influence of evolutionary pressures, such as mutation and selection. Despite its simple mathematical formulation, exact results for the distribution of allele frequency (DAF) as a function of time are not available in closed analytic form. Existing approximations build on the computationally intensive diffusion limit, or rely on matching moments of the DAF. One of the moment-based approximations relies on the beta distribution, which can accurately describe the DAF when the allele frequency is not close to the boundaries (zero and one). Nonetheless, under a Wright-Fisher model, the probability of being on the boundary can be positive, corresponding to the allele being either lost or fixed. Here, we introduce the beta with spikes, an extension of the beta approximation, which explicitly models the loss and fixation probabilities as two spikes at the boundaries. We show that the addition of spikes greatly improves the quality of the approximation. We additionally illustrate, using both simulated and real data, how the beta with spikes can be used for inference of divergence times between populations, with comparable performance to existing state-of-the-art method.

## INTRODUCTION

Advances in sequencing technologies have revolutionized the collection of genomic data, increasing both the volume and quality of available sequenced individuals from a large variety of populations and species (Romiguier *et al.* 2014; Gudbjartsson *et al.* 2015). These data, which may involve up to millions of single nucleotide polymorphisms (SNPs), contain information about the evolutionary history of the observed populations. There has been great focus in the recent years on inferring such histories and, to this end, one of the most widely used models is the Wright-Fisher (Gautier *et al.* 2010; Sirén *et al.* 2011; Malaspinas *et al.* 2012; Pickrell and Pritchard 2012; Gautier and Vitalis 2013; Steinrücken *et al.* 2014; Terhorst *et al.* 2015).

The Wright-Fisher model characterizes the evolution of a randomly mating population of finite size in discrete non-overlapping generations. The model describes the stochastic behavior in time of the number of copies (frequency) of alleles at a locus. The frequency is influenced by a series of factors, such as random genetic drift, mutations, migrations, selection, and changes in population size. When inferring the evolutionary history of a population, the effects of the different factors have to be untangled. The frequency varies from one generation to the next due to random sampling of a finite sized population (random genetic drift). Mutations, migrations and selection affect the sampling probability in a deterministic manner. We collectively refer to these as evolutionary pressures. Mutations and migrations result in linear changes of the sampling probability, while selection is a nonlinear pressure (Kimura 1964; Crow and Kimura 1970) and is therefore more difficult to study analytically.

A crucial step for carrying out statistical inference in the Wright-Fisher model is the determination of the distribution of the allele frequency (DAF) as a function of time, conditional on an initial frequency. Even though the Wright-Fisher model has a very simple mathematical formulation, no tractable analytical form exists for the DAF (Ewens 2004). Therefore, various approximations have been developed, ranging from purely analytical to purely numerical. They generally either build on the diffusion limit of the Wright-Fisher, or rely on matching moments of the true DAF. Both types of approximations have been used successfully for inference of populations divergence times (Sirén *et al.* 2011; Gautier and Vitalis 2013), populations admixture (Pickrell and Pritchard 2012), SNPs under selection (Gautier *et al.* 2010) and selection coefficients from time serial data (Malaspinas *et al.* 2012; Steinrücken *et al.* 2014; Terhorst *et al.* 2015).

Wright (1945) was the first to use the diffusion approximation to determine the stationary DAF. Kimura (1955) solved the diffusion limit and found the time-dependent distribution for pure drift, and Crow and Kimura (1956) extended the solution to include linear evolutionary pressures. However, these contain infinite sums, making their use cumbersome in practice. After decades dominated by inference based on the dual coalescent process (Rosenberg and Nordborg 2002; Hoban *et al.* 2012), diffusion has recently received increasing attention, and researchers have started to investigate other ways to solving analytically or approximating the diffusion equation (McKane and Waxman 2007; Waxman 2011; Malaspinas *et al.* 2012; Song and Steinrücken 2012; Zhao *et al.* 2013; Steinrücken *et al.* 2013; Steinrücken *et al.* 2014).

Moment-based approximations are less ambitious in that they aim at fitting mathematical convenient distributions by equating the first moments of the true DAF. Such approximations typically use either the normal distribution (Nicholson *et al.* 2002; Coop *et al.* 2010; Gautier *et al.* 2010; Pickrell and Pritchard 2012; Terhorst *et al.* 2015) or the beta distribution (Balding and Nichols 1995; Balding and Nichols 1997; Sirén *et al.* 2011; Sirén 2012). The rationale behind the use of these distributions is two-fold. Firstly, they are motivated by the diffusion limit: the normal distribution is the resulting DAF when drift is small (Nicholson *et al.* 2002), while the beta distribution is the stationary DAF under linear evolutionary pressures (Wright 1945; Crow and Kimura 1956). Secondly, they are entirely determined by their mean and variance. One major difference between the two is their support. Because the normal distribution is defined over the whole real line, it needs to be truncated to [0, 1] (Nicholson *et al.* 2002; Coop *et al.* 2010; Gautier *et al.* 2010). The truncated normal distribution has two atoms at zero and one (corresponding to the allele being lost or fixed) containing the densities in the intervals (*−∞,* 0] and [1, *∞*), respectively. However, the truncation procedure leads to a variance that no longer matches the variance of the true DAF (Gautier and Vitalis 2013). Alternatively, the full distribution can be applied for intermediary frequencies only, when the probabilities of lying outside the zero and one boundaries are small and can therefore be ignored (Pickrell and Pritchard 2012; Terhorst *et al.* 2015). Unlike the normal distribution, the beta distribution has the interval [0, 1] as support, but, due to its continuous nature, the probabilities at the boundaries will always be zero. Under a Wright-Fisher model, the loss and fixation events have a positive probability. The beta distribution provides a good fit for intermediary frequencies, but fails at capturing the non-zero boundary probabilities, as illustrated for pure drift in Figure 1A – C. When time is small, most of the probability mass is found close to the initial value *x*_0_ (Figure 1A). As time becomes larger, the allele frequency drifts away from *x*_0_ and more and more probability accumulates at the boundaries (Figure 1B and C).

**Figure 1:**
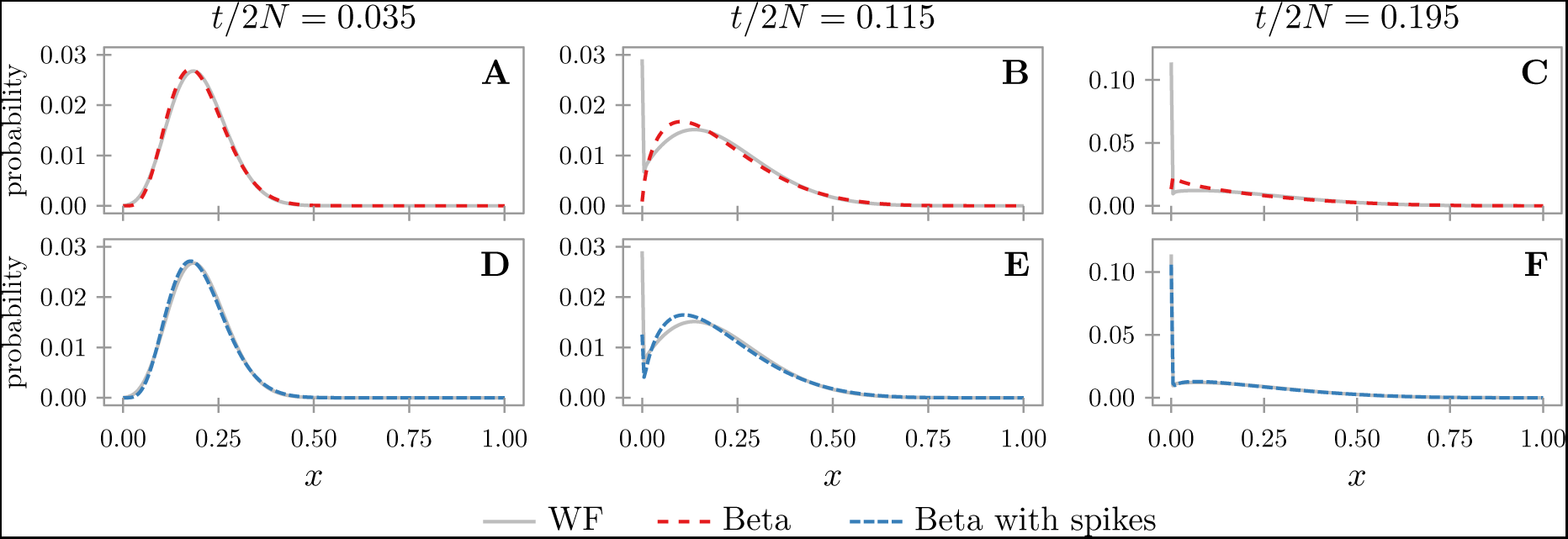
Fit of the beta and beta with spikes approximations. The figure shows the true discrete DAF as given by the Wright-Fisher model with a population size 2*N* = 200 under pure drift, and the corresponding discretized beta (A – C) and beta with spikes (D – F) approximations. The distributions are conditional on an initial frequency *x*_0_ = 0.2 and for different time points: *t/*2*N* = 0.035 (A, D), *t/*2*N* = 0.115 (B, E) and *t/*2*N* = 0.195 (C, F), where *t* is the number of discrete generations that the population has evolved. The discretization procedure is detailed in the Supplementary Material.

Here, we propose an accurate extension of the beta distribution under linear evolutionary pressures, entitled the beta with spikes, which explicitly models the probabilities at the boundaries. We show that the addition of spikes greatly improves the fit to the true DAF. We use simulation experiments and published chimpanzee exome data to demonstrate that the beta with spikes can be used for inference of population divergence times under pure drift, with performance comparable with a state-of-the-art diffusion-based method, and less computational burden. We additionally discuss how the beta with spikes can be used in future development to account for variable population size and selection.

Here, we propose an accurate extension of the beta distribution under linear evolutionary pressures, entitled the beta with spikes, which explicitly models the probabilities at the boundaries. We show that the addition of spikes greatly improves the fit to the true DAF. We use simulation experiments and published chimpanzee exome data to demonstrate that the beta with spikes can be used for inference of population divergence times under pure drift, with performance comparable with a state-of-the-art diffusion-based method, and less computational burden.

## THE BETA WITH SPIKES APPROXIMATION

Consider a diploid randomly mating population of size 2*N* and a biallelic locus with alleles *A*_1_ and *A*_2_. Under a Wright-Fisher model, the count of one of the alleles, *A*_1_, at the discrete generation *t* is a random variable *Z*_*t*_ *∈* {0, 1, *…,* 2*N}*. Let *X*_*t*_ = *Z*_*t*_/(2*N*) be the corresponding allele frequency. The evolution of *Z*_*t*_ is shaped by random genetic drift and deterministic evolutionary pressures. We capture the joint effect of the deterministic pressures in *g*(*x*), a polynomial in the allele frequency 0 *≤ x ≤* 1. Conditional on *Z*_*t*_, *Z*_*t*+1_ follows a binomial distribution (Ewens 2004)

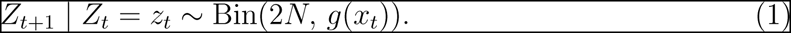

Here, we only consider linear evolutionary pressures, such as mutation and migration. Then *g*(*x*) takes the form

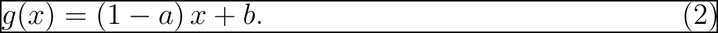

The parameters *a* and *b* verify that 0 *≤ b ≤ a ≤* 1 such that 0 *≤ g*(*x*) *≤* 1 for all 0 *≤ x ≤* 1. The case where *a* = 1, for which *g*(*x*) = *b* for all 0 *≤ x ≤* 1, has no biological meaning and we therefore assume that *a ≠* 1.

Under pure drift, *a* = *b* = 0. If mutations happen with probabilities *u* (from *A*_1_ to *A*_2_) and *v* (from *A*_2_ to *A*_1_), then *a* = *u* + *v* and *b* = *v*. Migration can be modeled, for example, by assuming that individuals can migrate away from the population under study and that there is an influx of individuals from a large population with constant frequency *x*_*c*_. Then, with probabilities *m*_1_ and *m*_2_, individuals migrate from and to the population under study, respectively. We have *a* = *m*_1_ and *b* = *m*_2_*x*_*c*_. Mutation and migration can be modeled jointly, resulting in *a* = *m*_1_ + (1 − *m*_1_)(*u* + *v*) and *b* = (1 − *m*_1_)*v* + *m*_2_*x*_*c*_. In the following, we treat the general linear case.

We are interested in the distribution of allele frequency (DAF) *X*_*t*_ conditional on *X*_0_ = *x*_0_, as a function of the generation *t*,

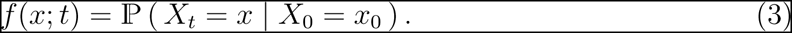

For simpler notation, we leave out the explicit condition on *X*_0_ = *x*_0_, and implicit condition on population size and evolutionary pressures.

Under the beta approximation, the DAF is

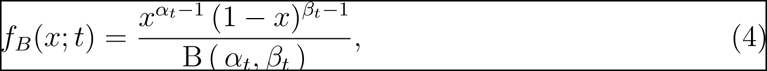

where B (*α, β*) is the beta function. The two shape parameters of the beta distribution are entirely determined by its mean and variance,

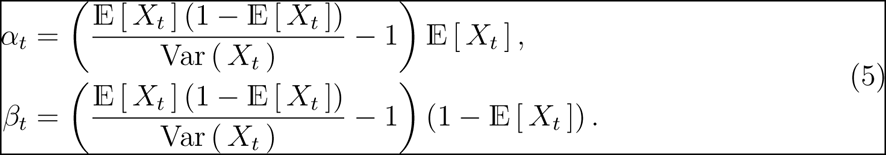

Therefore, in order to fit *f*_*B*_ to *f*, we need to calculate 𝔼 [*X*_*t*_] and Var (*X*_*t*_). These can be obtained in closed analytical form (see Supplementary Material for full derivation). The mean is entirely determined by the initial frequency *x*_0_ and the parameters *a* and *b* of the linear evolutionary pressures, while the variance also depends on the population size. Under pure drift (*a* = *b* = 0) we have

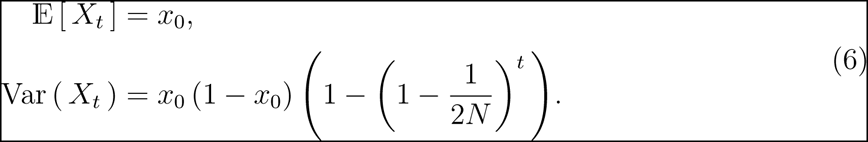

When *a* ≠ 0 we get

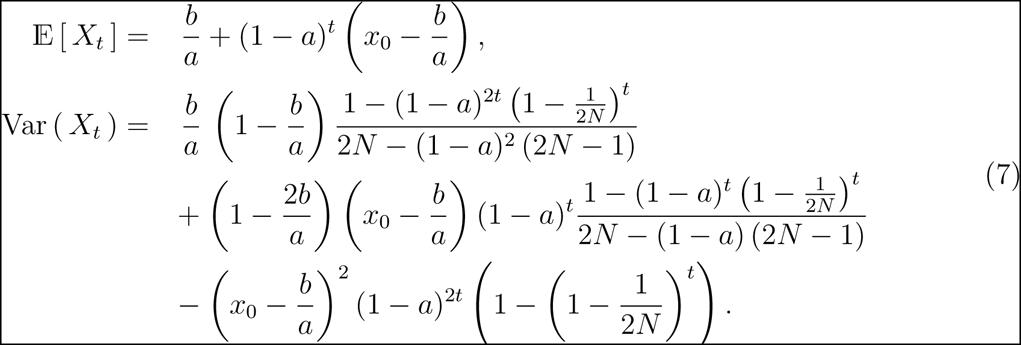

In the limit of infinite population size, the above formulas are equivalent to the mean and variance obtained by Sirén (2012) (up to some minor typographical errors, as confirmed by correspondence with the author; see also Supplementary Material).

To account for loss and fixation probabilities, we surround the beta distribution with two spikes

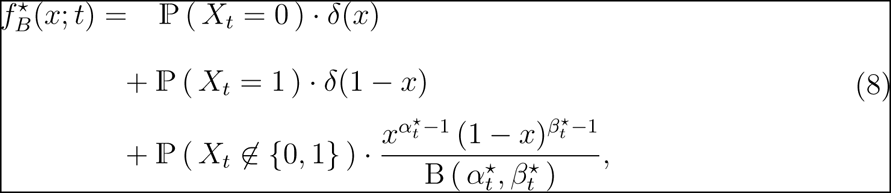

where *δ*(*x*) is the Dirac delta function and ℙ (*X*_*t*_ *∉*{0, 1}) = 1 − ℙ (*X*_*t*_ = 0) - ℙ (*X*_*t*_ = 1). To fit 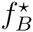to *f*, we need to determine the mean and variance of *X*_*t*_ conditional on polymerphism (*X*_*t*_ ∉ {0, 1*}*), and the probabilities ℙ (*X*_*t*_ = 0) and ℙ (*X*_*t*_ = 1) of loss and fixation, respectively. Given 𝔼 [ *X*_*t*_], Var (*X*_*t*_), ℙ (*X*_*t*_ = 0) and ℙ (*X*_*t*_ = 1), the conditional mean and variance can easily be calculated (see Supplementary Material). Therefore, we only require means of calculating the loss and fixation probabilities in order to fully specify the beta with spikes approximation. We use a recursive approach where we calculate the probabilities for *X*_*t*+1_ by relying on the approximated 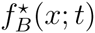 We additionally assume that *a* and *b* are small to obtain the following approximation for loss and fixation probabilities (see Supplementary Material for full derivation)

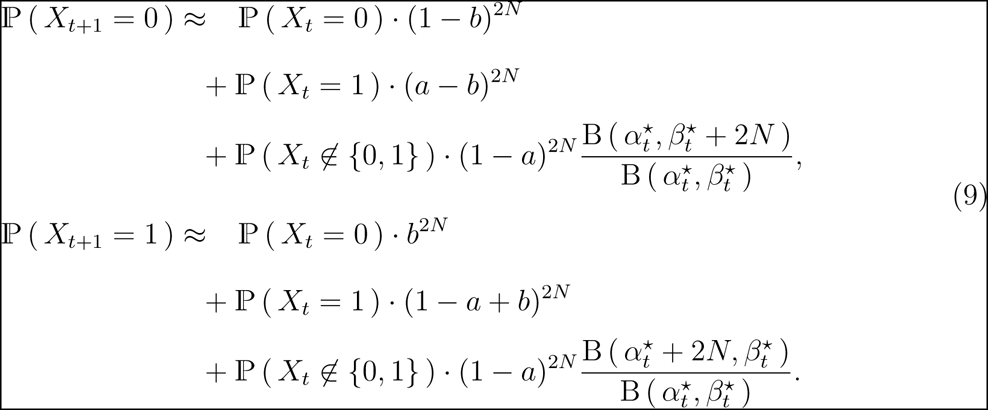

Figure 1D – F depicts the beta with spikes approximation for the same cases as in Figure 1A – C. When time is small (Figure 1A and D), the beta and beta with spikes distributions are equivalent, but as the time becomes larger, the advantage of adding the spikes becomes evident. As illustrated in supplementary Figure S1, the addition of spikes drastically improves the fit of the beta approximation to the true DAF under a Wright-Fisher model.

## INFERENCE OF DIVERGENCE TIMES

To further illustrate the advantage of incorporating the spikes, we inferred divergence times between populations, using both simulated data and exome sequencing data from three chimpanzee subspecies (Bataillon *et al.* 2015).

Populations are represented as successive descendants of a single ancestral population. We assume that after each split, the new populations evolved in isolation (no migration) under pure drift. A rooted tree (Figure 2) can be used to describe the joint history of several present populations, located at the leaves, while the common ancestral population is represented as the root. The data *𝔇* = *{*(*z*_*ij*_, *n*_*ij*_) *|* 1 *≤ i ≤ I,* 1 *≤ j ≤ J*} consist of *I* independent SNPs for *J* populations in the present: the (arbitrarily defined) reference (*A*_1_) allele count *z*_*ij*_ in a sample of size *n*_*ij*_ (0 *≤ z*_*ij*_ *≤ n*_*ij*_) for each locus 1 *≤ i ≤ I* and population 1 *≤ j ≤ J*.

**Figure 2:**
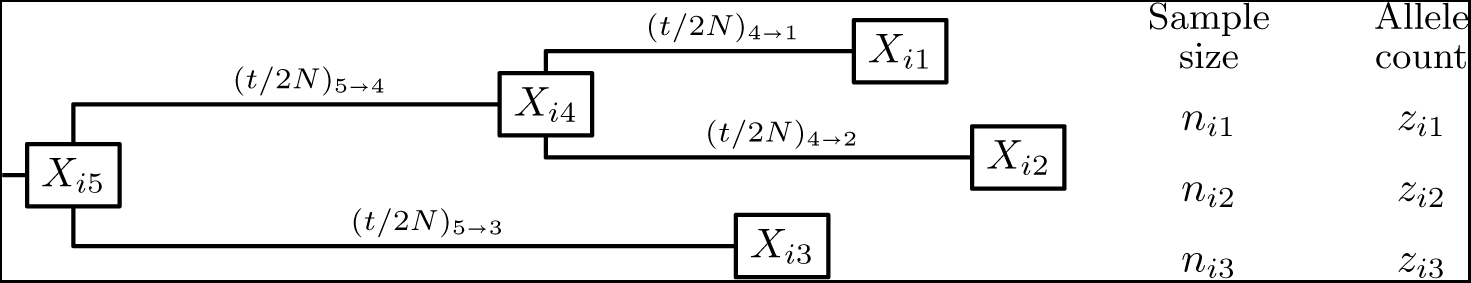
History of three populations in the present. The ancestral population 5 splits in populations 3 and 4, which further splits in populations 2 and 1. For each SNP *i* and present population *j* ∈ {1, 2, 3}, the data consists of the sample size *n*_*ij*_ and allele count *z*_*ij*_. The branch length between populations *k* and *j* is given as (*t/*2*N*)_*k⟶j*_ and represents the scaled number of generations that population *j* evolved since the split from the ancestral population *k*. The unknown allele frequencies of each population are denoted as *X*_*ij*_, with 1 *≤ j ≤* 5.

Conditional on the topology (i.e. tree without branch lengths), we inferred the scaled branch lengths by numerically maximizing the likelihood of the data.

### Likelihood of the data

Assuming Hardy-Weinberg equilibrium, the probability of observing *z*_*ij*_ alleles in a sample of size *n*_*ij*_ given the population allele frequency *x*_*ij*_ is given by the binomial distribution

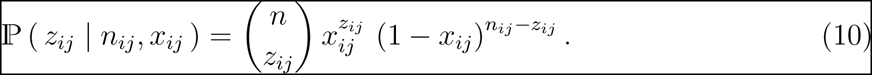

However, the allele frequencies *x*_*ij*_ are unobserved and the likelihood of the data 𝔇_*i*_ for SNP *i* is obtained by integrating over the unknown allele frequencies

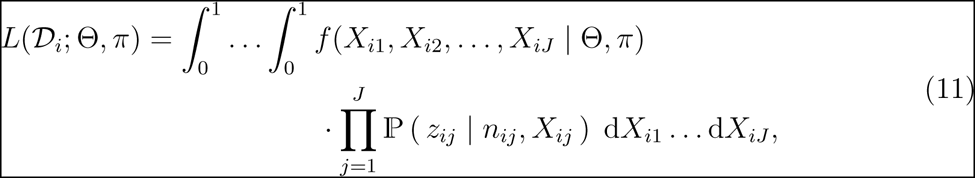

where *f* (*X*_*i*1_, *X*_*i*2_, …, *X*_*iJ*_ | Θ, *π*) is the joint distribution of the *X*_*ij*_’s at the leaves. The likelihood is a function of the scaled branch lengths, denoted here as Θ, and *π*, the unknown DAF at the root. The joint distribution *f* (*X*_*i*1_, *X*_*i*2_, …, *X*_*iJ*_ | Θ, *π*) is, in turn, an integral over the allele frequencies in the ancestral populations, represented as internal nodes in the tree. We approximate the integrals with sums by discretizing the allele frequencies. The discretized joint distribution is then obtained using a peeling algorithm (Felsenstein 1981), where the transition probabilities on each branch are given by the DAF (see Supplementary Material for details). As the SNPs are assumed to be independent, the full likelihood is a product over the SNPs,

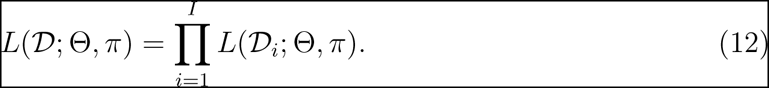

As SNP data contains only polymorphic sites, we further condition the above likelihood on polymorphic data as follows

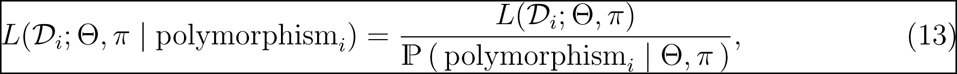

Where

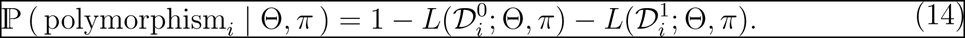

Here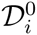 and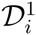 are data corresponding to site *i* where the allele was lost or fixed, respectively, in the samples from all populations,

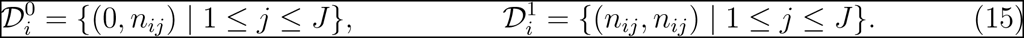

We treat *π*, the root DAF, as a nuisance parameter assumed to be a beta distribution. For a given topology (i.e. tree without branch lengths), the most likely branch lengths and shape parameters of *π* can be recovered by numerically maximizing the likelihood conditional on polymorphism.

### Simulated data

Using the topology depicted in Figure 2, we simulated multiple data sets containing independent SNPs under a Wright-Fisher model, given an acenstral frequency *X*_*i*5_ sampled from *π*, the root DAF, which we set to be a beta distribution. We used two different scenarios, labeled I and II, summarized in Table 1. Scenario I has a uniform *π* and large sample sizes, while scenario II is built to produce data that resembles the chimpanzee exome data analyzed below. For this, we used the chimpanzee sample sizes, and scaled branch lengths and root DAF as inferred by the beta with spikes on the chimpanzee data (see also Table 2).

**Table 1:** Simulation study scenarios.

|  | scenario I | scenario II |
| --- | --- | --- |
| $(t/2N)_{4 \rightarrow 1}$ | $40/(2 \cdot 200) = 0.1$ | $132/(2 \cdot 500) = 0.132$ |
| $(t/2N)_{4 \rightarrow 2}$ | $40/(2 \cdot 150) = 0.133$ | $44/(2 \cdot 500) = 0.044$ |
| $(t/2N)_{5 \rightarrow 4}$ | $40/(2 \cdot 150) = 0.133$ | $14/(2 \cdot 250) = 0.028$ |
| $(t/2N)_{5 \rightarrow 3}$ | $80/(2 \cdot 200) = 0.2$ | $300/(2 \cdot 250) = 0.6$ |
| shape parameters of $\pi$ | 1, 1 | 0.0188, 0.0195 |
| number of SNPs | 5,000 | 10,000 |
| sample sizes $n_{i1}, n_{i2}, n_{i3}$ | 100, 100, 100 | 22, 24, 12 |
| replicates | 50 | 50 |

The table indicates the values used for the branch lengths ( $t$ ), population sizes ( $N$ ) and scaled branch lengths ( $t/2N$ ), shape parameters of the beta distribution $\pi$ , the root DAF, number of SNPs and sample sizes used in the two simulation scenarios.

**Table 2:**
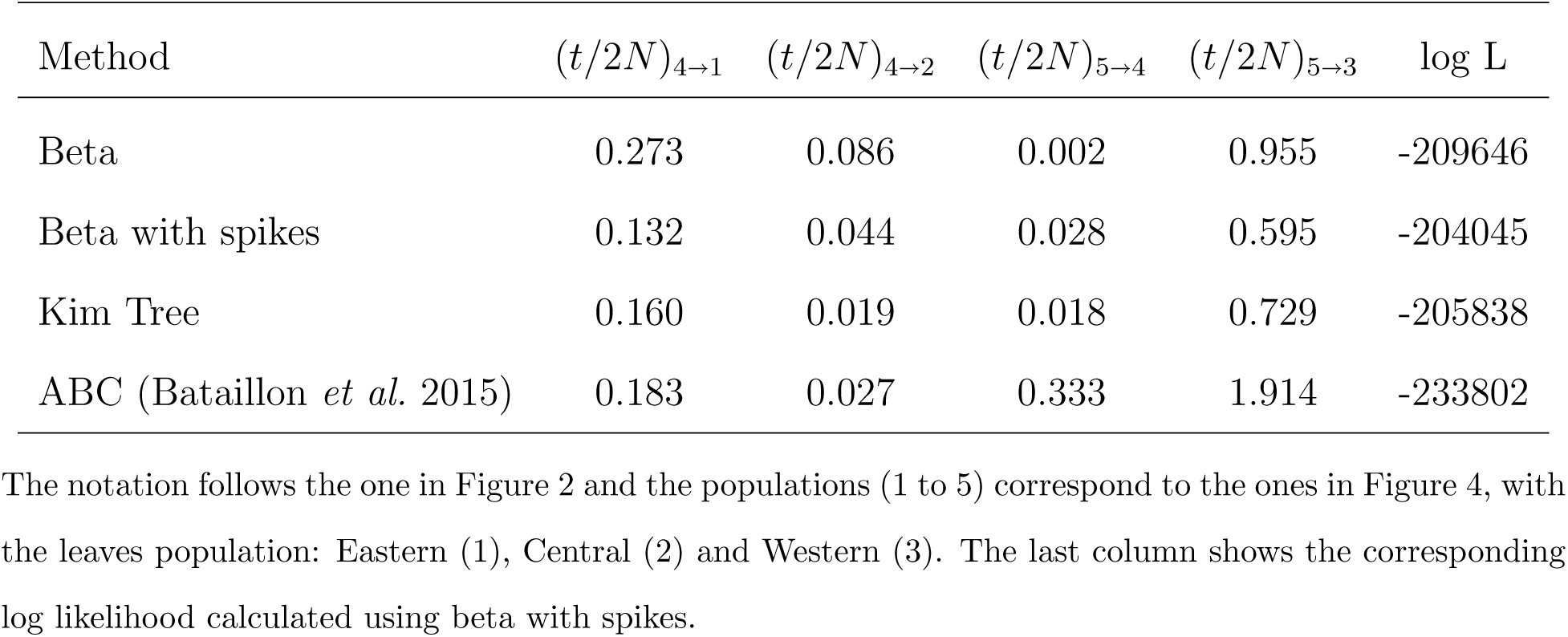
Inferred scaled branch lengths for the chimpanzee exome data.

| Method | $(t/2N)_{4 \rightarrow 1}$ | $(t/2N)_{4 \rightarrow 2}$ | $(t/2N)_{5 \rightarrow 4}$ | $(t/2N)_{5 \rightarrow 3}$ | log L |
| --- | --- | --- | --- | --- | --- |
| Beta | 0.273 | 0.086 | 0.002 | 0.955 | -209646 |
| Beta with spikes | 0.132 | 0.044 | 0.028 | 0.595 | -204045 |
| Kim Tree | 0.160 | 0.019 | 0.018 | 0.729 | -205838 |
| ABC (Bataillon *et al.* 2015) | 0.183 | 0.027 | 0.333 | 1.914 | -233802 |

For each simulated data set, we estimated the branch lengths using both the beta and beta with spikes as described previously. We additionally ran Kim Tree (Gautier and Vitalis 2013) using the default settings. Kim Tree is a method designed for inference of divergence times between populations evolving under pure drift. It uses Kimura’s solution to the diffusion limit for the DAF (Kimura 1955) and relies on a Bayesian MCMC approach. Here, we use the posterior means of the branch lengths as point estimates.

All methods estimate well the branches leading to populations 1 and 2 (Figure 3). Beta with spikes estimates the branch lengths more accurately and with lower variance than the beta approximation (see also supplementary Figure S2). Despite the fact that the spikes probabilities do not perfectly match the true loss and fixation probabilities (Figure 1E and F), this seems to have little effect on the accuracy of branch length estimation for beta with spikes. For both scenarios, the branch leading to population 2 and the inner branch from the root to population 4 have similar lengths, but the beta approximation and Kim Tree provide a worse estimate for the inner branch. This could be due to the fact that there is no data available resulting directly from the evolution on that branch, making the estimation problem harder. A similar result was obtained by Gautier and Vitalis (2013), where trees with the same topology were used. Interestingly, beta with spikes recovers the inner branch much more accurately than either beta and Kim Tree. When measuring the accuracy of the inferred lengths as an average over all four branches (supplementary Table S1), it is clear that beta with spikes outperforms Kim Tree for both scenarios.

**Figure 3:**
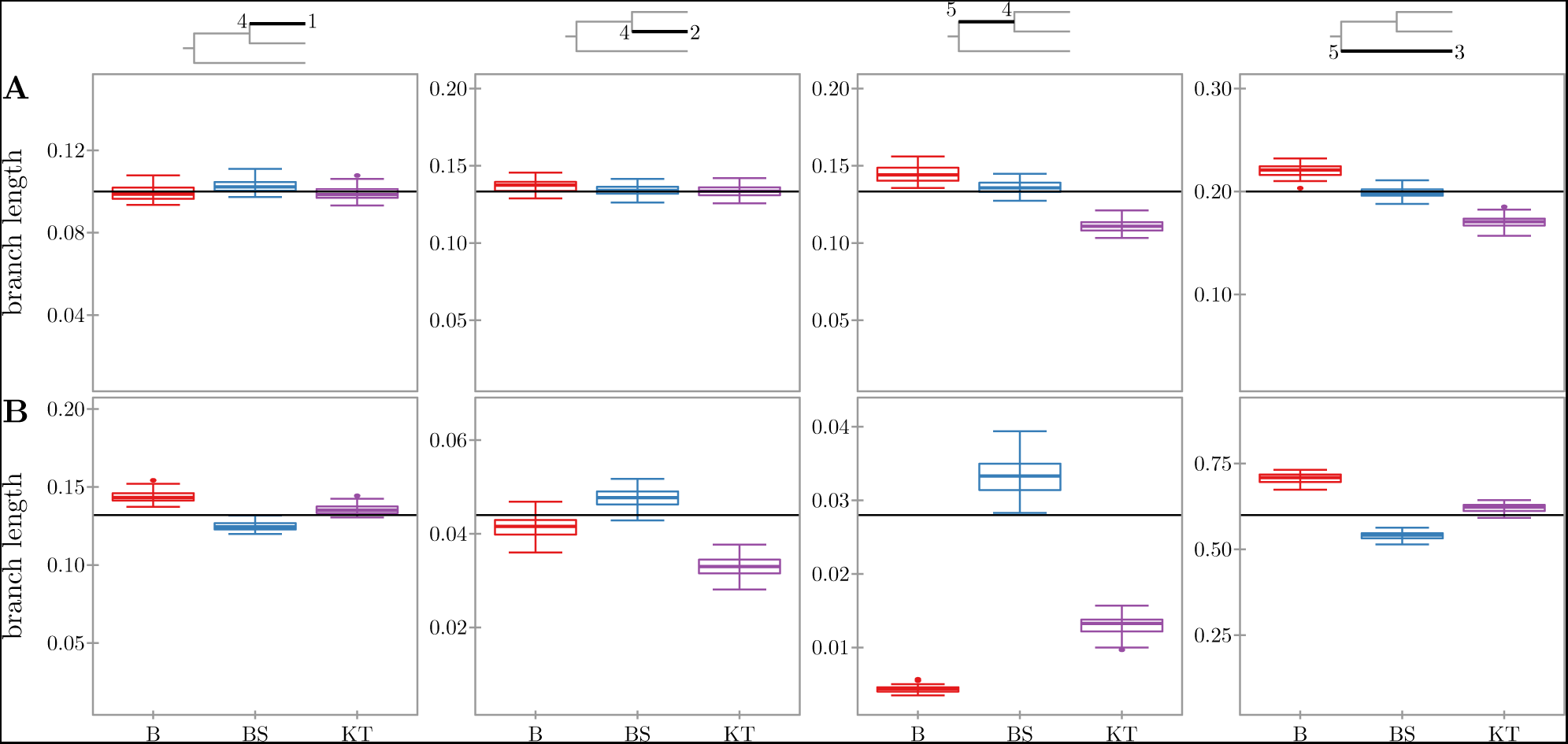
Inference of divergence times for simulation scenarios I (A) and II (B). The figure shows boxplots summarizing the inferred lengths for the four branches of the tree, indicated at the top of each column in black. The inferred lengths are plotted for beta (B), beta with spikes (BS) and Kim Tree (Gautier and Vitalis 2013) (KT). The (true) simulated length of each branch is plotted as a horizontal line. Each plot is scaled relative to the corresponding simulated branch length *τ*, with the limits of the y-axis set to [*τ* · 0.1, *τ* · 1.5].

### Chimpanzee data

The chimpanzee data analyzed here consisted of allele counts of autosomal synonymous SNPs obtained from exome sequencing of the Eastern, Central and Western chimpanzee subspecies (Bataillon *et al.* 2015) for 11, 12 and 6 individuals, respectively. From the original data set containing 59,905 synonymous SNPs, we filtered the SNPs where there was missing data, obtaining a total of 42,063 SNPs. We inferred the scaled branch lengths (Figure 4 and Table 2) using beta, beta with spikes and Kim Tree on the full data set and on 50 smaller data sets containing only 10,000 randomly sampled SNPs. Beta with spikes and Kim Tree infer comparable branch lengths, with the exception of the branch leading to the Western chimpanzee subspecies (population 3). We additionally report in Table 2 the likelihood of the full data calculated using beta with spikes for the branch lengths inferred using the three methods and the ones reported in the original study (Bataillon *et al.* 2015). Bataillon *et al.* (2015) used an ABC approach to fit a demographic model to the synonymous SNPs. Their results are consistent with the ones obtained here for the branches leading to the Eastern (population 1) and Central (population 2) chimpanzees. However, we obtain very different estimates for the remaining two branches. The likelihood in Table 2 indicates that the differences between beta with spikes and beta / Kim Tree / ABC are not merely a result of the numerical optimization being trapped in a local optimum, as the branch lengths obtained by the beta with spikes have the highest likelihood. The discrepancy between the beta with spikes and the ABC results is, perhaps, not surprising, as the difference in inferred branch lengths seems to correlate with the goodness of fit of the ABC demographic model to the observed data. Bataillon *et al.* (2015) report that their inferred demographic model shows a very good fit for the Central chimpanzees (difference in inferred branch length: 0.017), a slightly less good fit for the Eastern chimpanzees (difference in inferred branch length: 0.051) and a poorer fit for the Western chimpanzees (difference in inferred branch length: 1.319).

**Figure 4:**
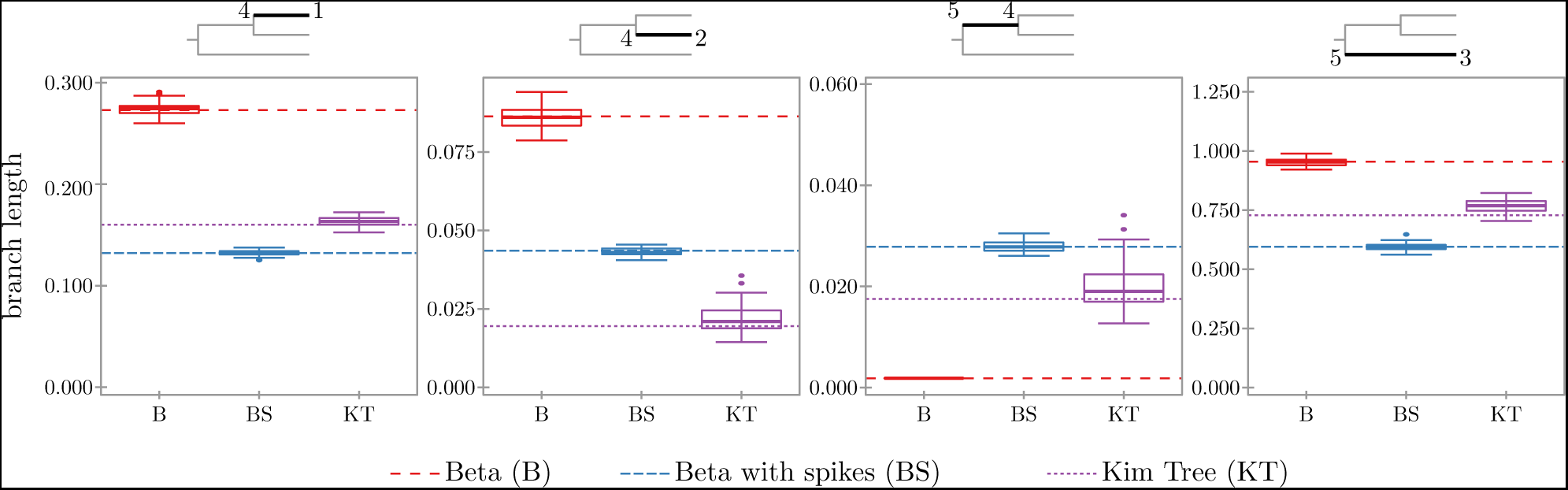
Inference of divergence times for the chimpanzee exome data. The figure shows boxplots summarizing the inferred lengths using 50 data sets with 10,000 SNPs that where randomly sampled from the full data set. The corresponding tree branches are indicated at the top of each plot in black. The inferred lengths are plotted for beta (B), beta with spikes (BS) and Kim Tree (Gautier and Vitalis 2013) (KT). The non-solid lines indicate the inferred lengths when running the methods on the full data set of 42,064 SNPs. The populations at the leaves are: Eastern (1), Central (2) and Western (3). Each plot is scaled relative to the corresponding branch length *τ* inferred by the beta with spikes on the full data set. The limits of the y-axis are set to [*τ ·* 0.05, *τ ·* 1.5].

## DISCUSSION

We have developed a new approximation to the distribution of allele frequency (DAF) as a function of time, conditional on an initial frequency, under a Wright-Fisher model with linear evolutionary pressures. Our work provides an accurate extension of the beta approximation (Balding and Nichols 1995; Balding and Nichols 1997; Sirén *et al.* 2011; Sirén 2012). As noted by Gautier and Vitalis (2013), the beta distribution ignores the possibility of loss or fixation of alleles. We addressed this issue by explicitly modeling the loss and fixation probabilities as two spikes at the boundaries. We showed that the addition of the spikes improves the quality of the approximation and results in more exact inference of divergence times between populations. We expect the beta with spikes to provide a less accurate approximation to the true DAF than the diffusion limit. Nevertheless, we showed that it can infer divergence times just as accurately as Kim Tree (Gautier and Vitalis 2013), a software built for inference of divergence times using Kimura’s solution to the diffusion limit (Kimura 1955).

### Computational complexity

The advantage of the beta with spikes becomes more clear when one considers its computational complexity. Diffusion methods rely on heavy computations, such as calculations of Gegenbauer polynomials (Gautier and Vitalis 2013), spectral decomposition of large matrices (Steinrücken *et al.* 2013; Steinrücken *et al.* 2014) or matrix inverse (Zhao *et al.* 2013). In contrast, the beta with spikes requires operations which are performed in constant time per iteration. Perhaps the most expensive evaluation is the beta function used in the loss and fixation probabilities, but very efficient approximations exist for this (Abramowitz and Stegun 1964). The difference in computational complexity is noticeable when comparing the running times of beta with spikes, implemented in `python 2.7`, and Kim Tree, implemented in `Fortran`. For the chimpanzee data set of 42,063 SNPs, beta with spikes ran in just under 5 minutes, while Kim Tree took almost an hour, even though `python 2.7` is a programming language less efficient than `Fortran`. We also note that the two inference methods are inherently different, as here we used a numerical optimization procedure, while Kim Tree uses a Bayesian MCMC approach.

### Extensions

We end this section by discussing possible extensions of the beta with spikes approximation and how these can be used in inference problems. Throughout this paper, we assumed that the population size is constant. Due to its recursive formulation, the beta with spikes lends itself naturally to incorporating variable population size, without any increase in computational complexity. This can then be used for inference of population size backwards in time, similar to methods relying on the coalescent with recombination (Li and Durbin 2011; Sheehan *et al.* 2013; Schiffels and Durbin 2014). A recently published method (Liu and Fu 2015) illustrates that allele frequency data, summarized as site frequency spectra, can be efficiently used for inference of variable population size backwards in time. Even though Liu and Fu (2015) assume sites are independent and do not use linkage information, their method can handle larger data sets than Li and Durbin (2011), which leads to more accurate inference of population sizes for the recent past. The results obtained by Liu and Fu (2015) indicate that the beta with spikes could be successfully used for such demographic inference.

Another extension of the presented approximation would be to incorporate selection, which is a non-linear evolutionary pressure. In the recent years, there has been a great focus on inference of selection coefficients from time-series data under a Wright-Fisher model (Malaspinas *et al.* 2012; Bank *et al.* 2014; Steinrücken *et al.* 2014; Foll *et al.* 2015; Terhorst *et al.* 2015). A newly developed statistical method aims at modeling the evolution of multi-locus alleles under a Wright-Fisher model with selection (Terhorst *et al.* 2015), by fitting a multivariate normal distribution from the first moments of the DAF. Using the approach of Terhorst *et al.* (2015) for moment calculation, the beta with spikes can be extended to non-linear evolutionary pressures. Terhorst *et al.* (2015) do not treat the loss and fixation probabilities. However, as selection is expected to drive allele frequencies towards the boundaries faster than pure drift, including the explicit spikes becomes crucial.

## AVAILABILITY

The beta, beta with spikes approximations, inference of divergence times and simulation under a Wright-Fisher model were implemented in `python 2.7`. The code is freely available at https://github.com/paula-tataru/SpikeyTree.

## ACKNOWLEDGMENTS

It is a pleasure to thank Thomas Mailund for helpful discussions. This work has been supported, in part, by the European Research Council under the European Unions Seventh Framework Program (FP7/20072013, ERC grant number 311341) and the Danish Research Council (grant number DFF–4002-00382).

